# Gene fate spectrum as a reflection of local genomic properties

**DOI:** 10.1101/2022.08.08.502781

**Authors:** Yuichiro Hara, Shigehiro Kuraku

## Abstract

Functionally indispensable genes are likely to be retained and otherwise to be lost during evolution. This evolutionary fate of a gene can also be affected by neutral factors, including the mutability of genomic positions, but such features have not been examined well. To uncover the genomic features associated with gene loss, we investigated the characteristics of genomic regions where genes have been independently lost in multiple lineages. With a comprehensive scan of gene phylogenies of vertebrates with a careful inspection of evolutionary gene losses, we identified 1,081 human genes whose orthologs were lost in multiple mammalian lineages: designated ‘elusive genes.’ These elusive genes were located in genomic regions with rapid nucleotide substitution, high GC content, and high gene density. A comparison of the orthologous regions of such elusive genes across vertebrates revealed that these features had been established before the radiation of the extant vertebrates more than 500 million years ago. The association of human elusive genes with transcriptomic and epigenomic characteristics illuminated that the genomic regions containing such genes were subject to repressive transcriptional regulation. Thus, the heterogeneous genomic features driving gene fates toward loss have been in place since the ancestral vertebrates and may sometimes have relaxed the functional indispensability of such genes.

## Introduction

In the course of evolution, genomes continue to retain most genes with occasional duplications, while losing some genes. This retention and loss can be interpreted as gene fate; genes are stably retained in the genome, but some factors may cause them to transition to a state where deletion occurs. Accordingly, identification of the factors allowing gene loss may facilitate our understanding of gene fate. Gene retention or loss has generally been considered to depend largely on the functional importance of the particular gene from the perspective of molecular evolutionary biology (Albalat and Cañestro, 2016; Bartha et al., 2018; Blanc et al., 2012; Liu et al., 2015; Olson, 1999; Sharma et al., 2018; Shen et al., 2018). Genes with indispensable functions have usually been retained with highly conserved sequences in genomes, through rapid elimination of alleles that impair gene functions (Hirsh and Fraser, 2001; Krylov et al., 2003; Miyata et al., 1980; Pál et al., 2006). However, genes with less important functions are likely to accept more mutations and structural variations, which can degrade the original functions, leading to gene loss through pseudogenization or genomic deletion (Jordan et al., 2002; Yang et al., 2003). To date, gene loss has been imputed to the relaxation of functional constraints of individual genes. Gene loss has further been revealed to drive phenotypic adaptation in various organisms (Albalat and Cañestro, 2016; Olson, 1999), as well as in a gene knockout collection of yeasts in culture (Giaever and Nislow, 2014; Maclean et al., 2017).

To uncover the association between fates and functional importance of the genes, molecular evolutionary analyses have been conducted at various scales, from gene-by-gene to genome-wide. A number of studies have revealed that the genes with reduced non-synonymous substitution rates (or *K*_A_ values) and ratios of non- synonymous to synonymous substitution rates (*K*_A_/*K*_S_ ratios) are less likely to be lost (Jordan et al., 2002; Yang et al., 2003). A genome-wide comparison of duplicated genes in yeast revealed larger *K*_A_ values for those lost in multiple lineages than those retained by all the species investigated (Byrne and Wolfe, 2007). Other comprehensive studies of gene loss across metazoans and teleosts revealed that the genes expressed in the central nervous system are less prone to loss (Fernández and Gabaldón, 2020; Roux et al., 2017). These observations again suggest that gene fate depends on the functional constraints of a particular gene.

Besides functional constraints, several studies have identified the genes lost independently in multiple lineages, revealing that the genomic regions containing these genes ‘prefer’ particular characteristics associated with structural instability (Cortez et al., 2014; Hughes et al., 2012; Lewin et al., 2021; Maeso et al., 2016). In mammals, tandemly arrayed homeobox genes derived from the Crx gene family were lost in multiple species (Lewin et al., 2021; Maeso et al., 2016). The findings suggest that genomic features containing tandem duplications facilitate unequal crossing over, leading to frequent gene loss. Mammalian chromosome Y, which contains abundant repetitive elements and continues to reduce in size, has lost a considerable number of genes (Cortez et al., 2014; Hughes et al., 2012). Genes in such particular genomic regions may be prone to loss in a more neutral manner than the relaxation of functional importance or via functional adaptations. Accordingly, these studies focusing on the particular genomic regions led us to search for the common features in genomes that potentially facilitate gene loss. Genome-wide scans have revealed heterogeneous distributions of a variety of sequence and structural features so far, for example, base composition (Bernardi and Bernardi, 1986; Cohen et al., 2005; Katzman et al., 2011), the frequency of repetitive elements (Korenberg and Rykowski, 1988; Medstrand et al., 2002), and DNA-damage sensitivity induced by replication inhibitors (Debatisse et al., 2012; Helmrich et al., 2006). However, the extent to which these characteristics are associated with gene fates has not been understood well at a genome-wide level.

The accumulation of near-complete genome assemblies for various organisms facilitates comprehensive taxon-wide analysis of gene loss (Fernández and Gabaldón, 2020; Guijarro-Clarke et al., 2020; Rice and McLysaght, 2017). Along with this motivation, we recently performed a comprehensive analysis on the fate of paralogs generated via the two-round whole genome duplications in early vertebrates (Hara et al., 2018a). The results revealed that the genes retained by reptiles but lost in mammals and Aves rapidly accumulated not only non-synonymous but also synonymous substitutions in comparison with the counterparts retained by almost all the vertebrates examined, indicating that those genes prone to loss harbor rapid mutation rates. Furthermore, these loss-prone genes were located in genomic regions with high GC-contents, high gene densities, and high repetitive element frequencies. These findings suggest that the fates of those genes are influenced not only by functional constraints but also by intrinsic genomic characteristics. Because the findings were restricted to a set of particular genes, they prompted us to examine whether this trend is associated with gene fates on a genome-wide scale.

In this study, we inferred molecular phylogenies of vertebrate orthologs to systematically search for the genes harboring different fates in the human genome. We referred to the loss-prone genes as ‘elusive’ genes that were retained by modern humans but were lost independently in multiple mammalian lineages. As a comparison of the elusive genes, we retrieved the ‘non-elusive’ genes that were retained by almost all of the mammalian species examined. We conducted a careful search for gene loss to reduce the false discovery rate, which is usually caused by incomplete sequence information (Botero-Castro et al., 2017; Deutekom et al., 2019). By comparing the genomic regions containing these genes, we uncovered genomic characteristics relevant to gene loss. We associated the elusive genes with a variety of findings from deep sequencing analyses of the human genome including transcriptomics, epigenomics, and genetic variations. These data assisted us to understand how intrinsic features of genomes—presumably unrelated to gene function, may affect gene fate, leading to loss by relaxing the functional importance of ‘elusive’ genes.

## Results

### Identification of human ‘elusive’ genes

We defined an ‘elusive’ gene as a human protein-coding gene that existed in the common mammalian ancestors but was lost independently in multiple mammalian lineages (Figure 1; see Methods for details). In our analysis, we searched for such genes by reconstructing phylogenetic trees of vertebrate orthologs and detecting gene loss events within the individual trees. To search for elusive genes, we paid close attention to distinguishing true evolutionary gene loss from falsely inferred gene loss caused by insufficient genome assembly, gene prediction, and orthologous clustering (Botero-Castro et al., 2017; Deutekom et al., 2019), as described below.

**Figure 1.**
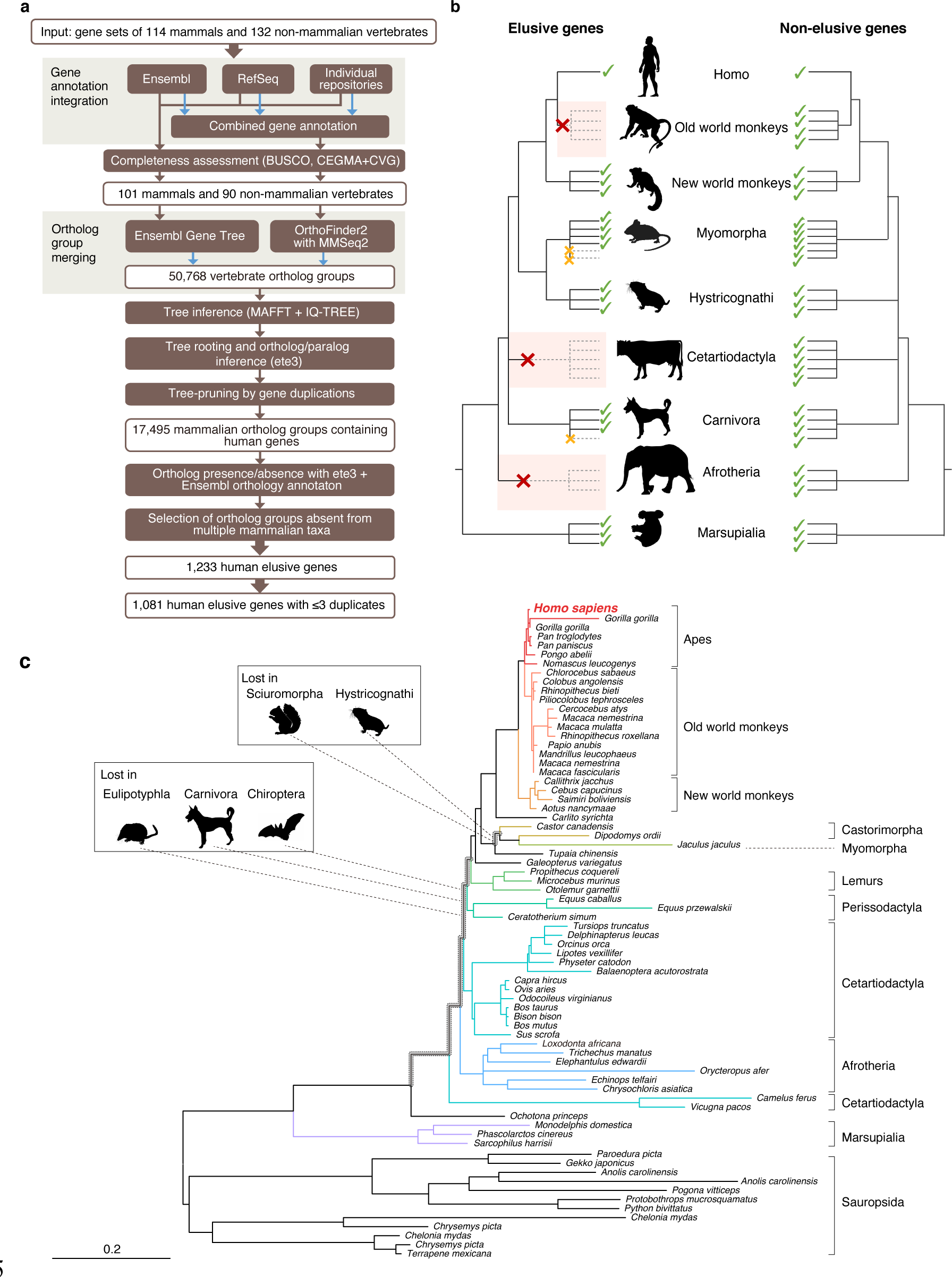
Detection of ‘elusive’ genes. (**a**) Pipeline of ortholog group clustering and gene loss detection. (**b**) Definition of an elusive gene schematized with ortholog presence/absence pattern referring to a ‘higher’ taxonomic hierarchy. (**c**) A representative phylogeny of the elusive gene encoding Chitinase 3-like 2 (CHI3L2). Taxa shown in the tree were used to investigate the presence or absence of orthologs. The Sciuromorpha, Hystricognathi, Eulipotyphla, Carnivora, and Chiroptera are absent from the tree, indicating that the CHI3L2 orthologs were lost somewhere along the branches framed in gray in the tree. In addition, the orthologs of many members of the Myomorpha were not found, suggesting that gene loss occurred in this lineage.

We first produced highly complete orthologous groups comprised of nearly complete gene sets. We merged multiple gene annotations of a single species followed by assessments of the completeness of the gene sets (Figure 1a). Using these gene sets, we then created two sets of ortholog groups with different methods and merged them into a single set (Figure 1a). In searching for gene loss events, we restricted our study to those that occurred in the common ancestors of particular ‘higher’ taxa. This procedure relieved false identifications of gene loss in a species or an ancestor of a lower taxonomic hierarchy caused by incomplete genomic information (Figure 1b).

We integrated gene annotations from Ensembl, RefSeq, and the sequence repositories of individual genome sequencing projects to produce gene annotations for 114 mammalian and 132 non-mammalian vertebrates. From these, we selected the annotations of 101 and 90 species, respectively, that exhibited high completeness in the BUSCO assessment (Simão et al., 2015) (Supplementary Table S1). Using these gene sets, ortholog clustering was conducted by OrthoFinder, and these ortholog groups were integrated into the ones provided by the Ensembl Gene Tree. This integration resulted in 50,768 vertebrate ortholog groups. Phylogenetic tree inference of the integrated ortholog groups and pruning of the individual trees based on gene duplications resulted in 17,495 mammalian ortholog groups that contained human genes. For the individual mammalian ortholog groups, we searched for family or ‘higher’ taxonomic groups (listed in Supplementary Table S1) in which the gene was absent in all the species examined (Figure 1b). We interpreted this gene absence as an evolutionary loss that occurred in the common ancestor of the taxon. Finally, we extracted the ortholog groups that were retained by humans but were lost independently in the common ancestors of at least two taxa (Figure 1c). Hereafter we call the human genes belonging to these ortholog groups ‘elusive genes.’ To compare these, we also selected the ortholog groups that contained all of the mammals examined including single-copy human genes. We called these ‘non-elusive genes.’ This comprehensive scan of gene phylogenies resulted in 1,081 elusive and 8,050 non-elusive genes (Supplementary Table S2).

### Genomic signatures of the human elusive genes

The loss-prone nature of the elusive genes suggests a relaxation of their functional constraints. To uncover the molecular evolutionary characteristics associated with each elusive gene, we computed synonymous and non-synonymous substitution rates, namely *K*_S_ and *K*_A_ values, respectively, between human and chimpanzee and mouse orthologs for the elusive and non-elusive genes. The results showed larger *K*_A_ values in the ortholog pairs of the elusive genes than in those of the non-elusive genes (Figure 2a; Figure 2–figure supplement 1). This indicates rapid accumulation of amino acid substitutions in the elusive genes, potentially accompanied by the relaxation of functional constraints. Our analysis further illuminated larger *K*_S_ values for the elusive genes than in the non-elusive genes (Figure 2b; Figure 2–figure supplement 1).

**Figure 2.**
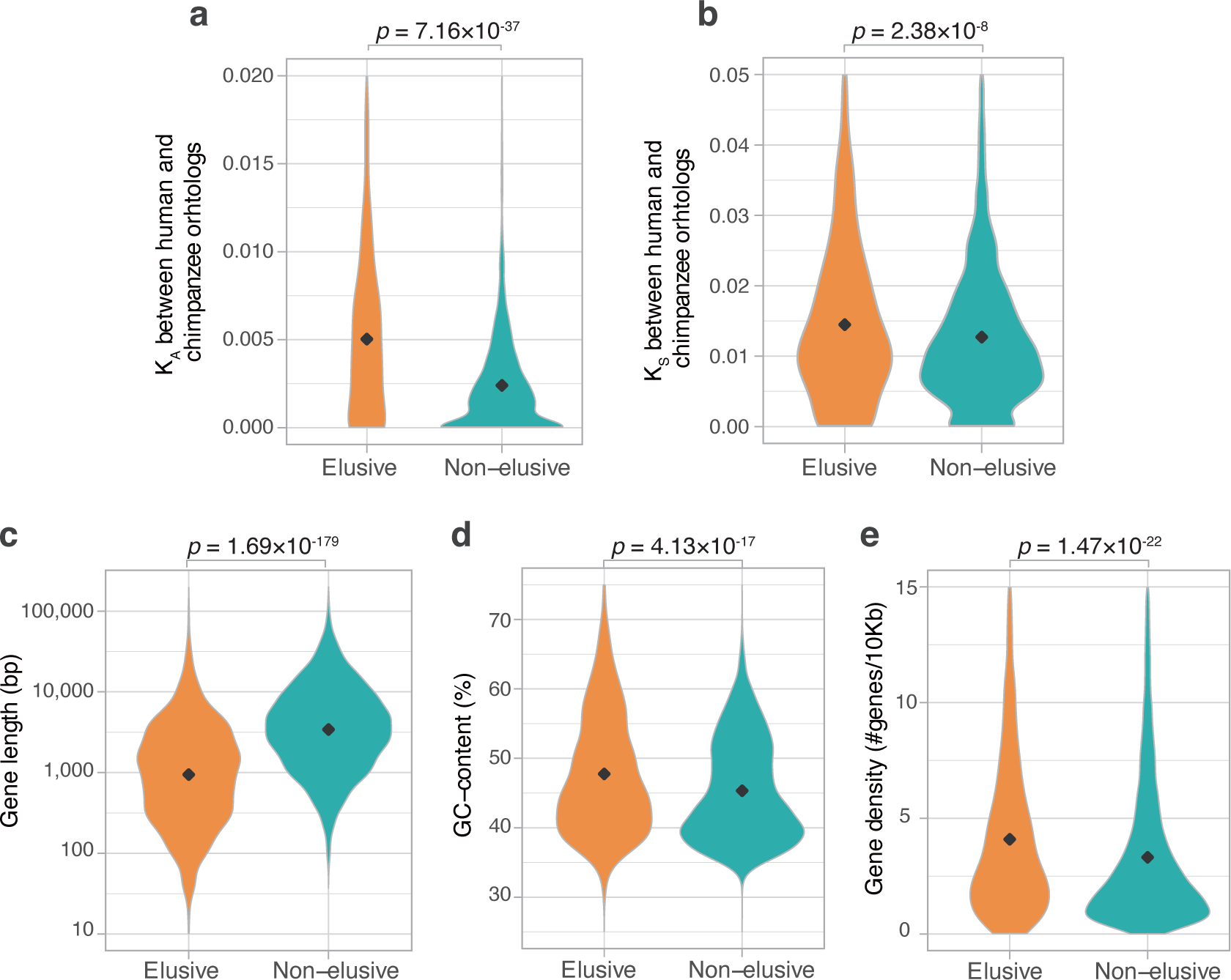
Genomic and evolutionary characteristics of elusive genes Distributions of non-synonymous and synonymous substitution rates, namely *K*_A_. (**a**) and *K*_S_ (**b**) values, respectively, between the human-chimpanzee orthologs of the elusive and non-elusive genes. Distribution of gene length (**c**) and GC content (**d**) of the human elusive and non-elusive genes. (**e**) Distribution of gene density in the genomic regions where the human elusive and non-elusive genes are located.

Importantly, the abundance of synonymous substitutions, which do not affect changes in amino acid residues, indicates that the elusive genes are also susceptible to genomic characteristics independent of selective constraints on gene functions.

To further scrutinize the characteristics reflecting the genomic environment rather than gene function, we analyzed genomic characteristics that may distinguish the elusive from non-elusive genes. A comparison between these two categories revealed shorter gene-body lengths and higher GC contents of elusive rather than non-elusive genes (Figure 2c,d). Furthermore, a scan of intergenomic gene distribution revealed that the elusive genes were located in the genomic regions with high gene density compared with the non-elusive genes (Figure 2e). Our findings indicate that such elusive genes have distinct characteristics in the human genome. These genomic characteristics, as well as high nucleotide substitution rates, were consistent with the findings in our genome analyses using the amniote and elasmobranch genomes (Hara et al., 2018b, 2018a).

### Tracing elusiveness back along the vertebrate evolutionary tree

The origins of the human elusive genes can be traced back along the evolutionary tree, at least to the mammalian common ancestor. To investigate possible antiquities of the genomic properties associated with elusive genes, we investigated their orthologs in non-mammalian vertebrate genomes. By scrutinizing the ortholog groups that were used for elusive gene identification, we identified 982 human elusive gene orthologs for chimpanzee, 540 for mouse, 380 for chicken, 415 for central bearded dragon, 416 for clawed frog, 415 for coelacanth, 431 for spotted gar, and 390 for bamboo shark. These four non-mammalian vertebrates retained orthologs of fewer than half of the elusive genes, but most of the non-elusive ones (Figure 3–figure supplement 1a). In the coelacanth, gar, and shark, the orthologs of the elusive genes were less frequently retained by all the species than those of the non-elusive ones (Figure 3–figure supplement 1b). This suggests that the origins of the loss-prone propensity of the elusive genes potentially date back to long before the emergence of the Mammalia.

**Figure 3.**
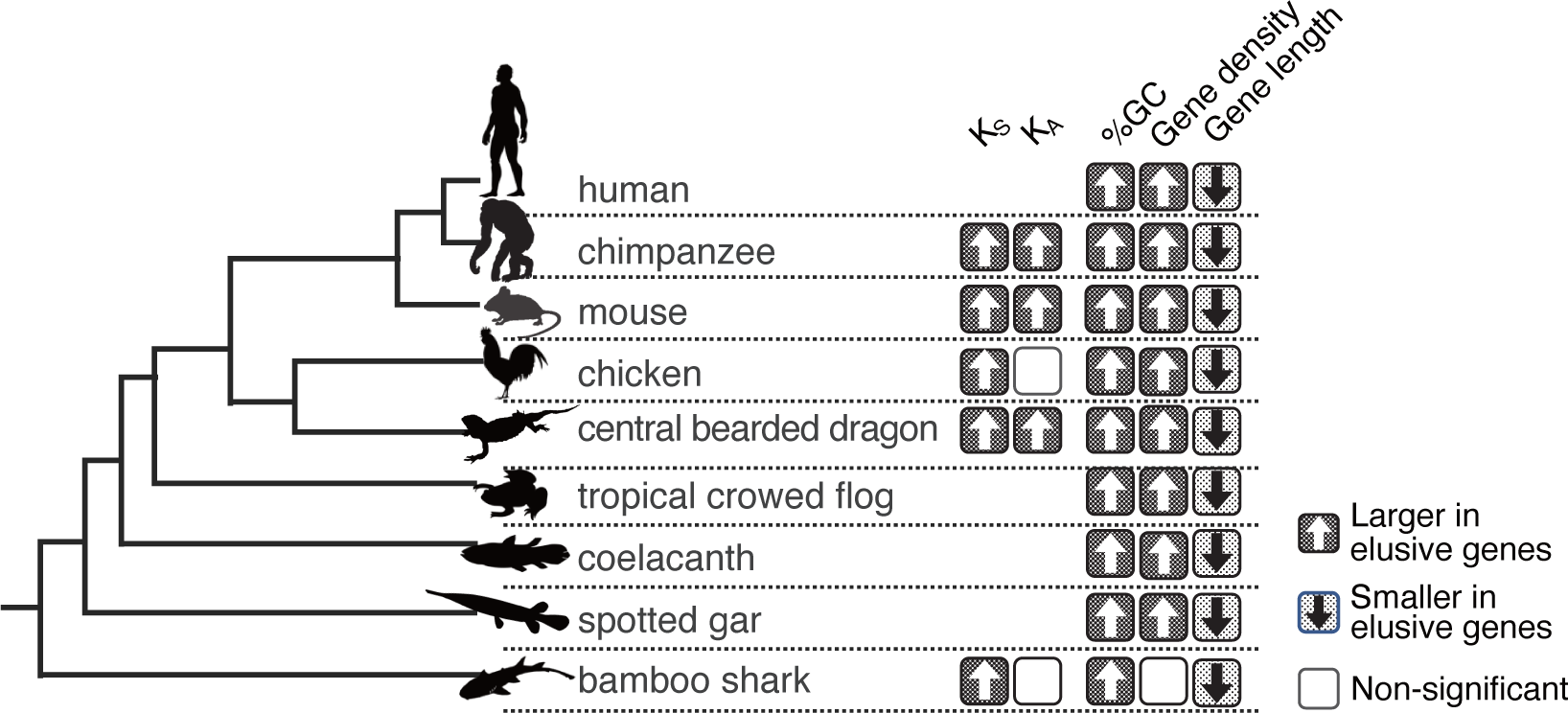
Longstanding characteristics of elusive genes. Retention of the genomic and evolutionary characteristics of the human elusive genes across vertebrates. The individual round squares with arrowheads indicate significant increases or decreases of the distribution of particular characteristics in the orthologs of the human elusive genes and their flanking regions, compared with those of the non- elusive genes in these selected vertebrate genomes. For the chimpanzee and mouse genomes, *K*_A_ and *K*_S_ values were computed between the human elusive genes and the orthologs of these mammals. For the non-mammalian species, these values were computed with ortholog pairs for the elusive/non-elusive genes between the corresponding species and their closely related species: turkey for chicken, green anole for central bearded dragon, and whale shark for bamboo shark. Distributions of these metrics for non-human species are shown in Supplementary Figures S1 and S3.

We further examined the genomic characteristics harbored by the human elusive genes in the vertebrate orthologs. In all the species examined, the orthologs of the elusive genes exhibited high GC content and compact gene bodies. Additionally, in most of these species, the orthologs of elusive genes were located in genomic regions with high gene density compared with orthologs of the non-elusive genes (Figure 3; Figure 3–figure supplement 2). In addition, we computed *K*_S_ and *K*_A_ values between the orthologs of the vertebrate species and their close relatives for elusive and non-elusive genes. In any of the species pairs, the orthologs of the elusive genes were found to harbor higher *K*_S_ values than those of the non-elusive gene orthologs, while the orthologs of the elusive genes exhibited higher *K*_A_ values in mammals and lizards (Figure 3; Figure 2–figure supplement 1). These observations indicate that these genomic characteristics probably originated before the emergence of gnathostomes, a monophyletic group of chondrichthyan and bony vertebrates, and have been retained for at least 500 million years.

### Abundant polymorphism in elusive genes

The observation of large *K*_S_ and *K*_A_ values in the elusive genes prompted us to examine the extent to which these genes have accommodated genetic variations in modern humans. Large-scale human genome resequencing projects have identified a huge number of genetic variations, from rare to common, and from single nucleotide variants (SNVs) to chromosome-scale structural variants, facilitating tackling this issue. We retrieved copy number variants (CNVs) and rare SNVs in the human genome from the Database of Genomic Variants, release 2016-08-31 (MacDonald et al., 2014) and dbSNP release 147 (Sherry et al., 2001), respectively, and computed their densities in the individual genic regions. We found that the genic regions of the human elusive genes contained abundant rare SNVs, as well as deletion and duplication CNVs, compared with those of the non-elusive genes (Figure 4a–c). This result suggests that genomic regions containing the elusive genes are not only prone to loss but also to duplication.

**Figure 4.**
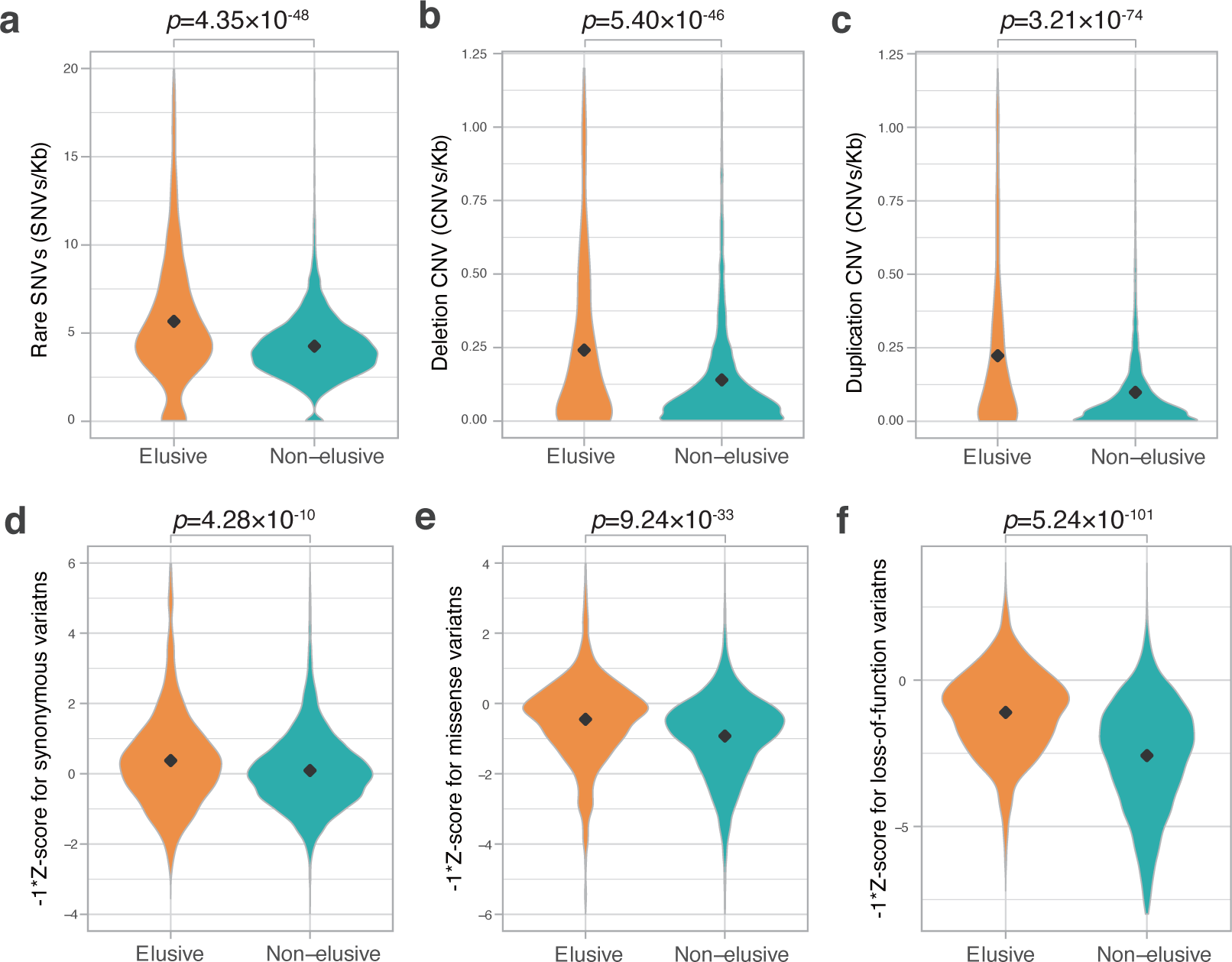
Genetic variations of the elusive and non-elusive genes within human populations Comparison of the density of rare SNVs. (**a**), deletion CNVs (**b**), duplication CNVs (**c**), and Z-scores of synonymous (**d**), missense (**e**), and loss-of-function variants (**f**). We used opposite numbers of the *Z*-scores in **d–f** so that the elusive genes have higher values than non-elusive genes as in Figures 2a,b, d, e and 3a–c.

To evaluate the functional consequences of abundant genetic variants in the elusive genes, we investigated genetic variations stored in the gnomAD v. 2.1 database, a repository containing >120,000 exome and >15,000 whole genome sequences of human individuals (Karczewski et al., 2021). This database classifies SNVs in coding regions into three categories—synonymous, missense, and loss-of-function—and the loss-of- function category contains nonsense mutations, frameshift mutations, and mutations in splicing junctions. The gnomAD site computes a *Z*-score, an index representing the abundance of SNVs for individual genes; positive and negative values denote fewer or more mutations in a coding region than expected, respectively (Figure 4d–f).

Accordingly, the *Z*-score for nonsense mutations and loss-of-function mutations of the individual genes indicates the degree of natural selection: larger values demonstrate genes subjected to purifying selection, while smaller ones suggest functional relaxation. We found lower *Z*-scores of missense and loss-of-function mutations (higher opposite numbers of *Z*-scores in Figure 4e, f) in the human elusive genes than in the non-elusive genes, suggesting that the elusive genes are more functionally dispensable and potentially resistant to harmful mutations. Additionally, the *Z*-scores of synonymous mutations of the human elusive genes were higher than those of the non-elusive genes (Figure 4d). This confirms the high mutability of genomic regions containing elusive genes, as observed in the *K*_S_ values.

### Transcriptomic natures of elusive genes

To further investigate how the human elusive genes have decreased functional essentiality, we examined their expression profiles. For this purpose, we compared gene expression profiles of the 54 adult tissues from the GTEx database v. 8 (GTEx Consortium, 2020) between the elusive and non-elusive genes. For individual genes, we computed the maximum Transcription Per Million (TPM) values among these tissues as the expression quantity level. For expression diversities, we employed Shannon’s diversity index *H*′, which is often utilized as an index of species diversity in the ecological literature, based on the proportion of TPM values across the 54 tissues.

As shown in the density scatter plots of the individual genes displaying these two indicators in Figure 5, most of the non-elusive genes possessed large maximum TPM and *H*′ values. Thus, most non-elusive genes are ubiquitously expressed at certain levels. By contrast, the density plot of the elusive genes displayed an additional high-density spot with small TPM and *H*′ values, indicating that the genes in this spot were not expressed, at least in adult tissues. The plot also showed another broad dense area of small *H*′ values, which contained the genes expressed in a single or a few tissues. A similar analysis was performed with the fetal single cell RNA-seq data (Cao et al., 2020), revealing that the averaged expression profiles of the elusive and non-elusive genes for the 172 cell types were concordant with those of the adult tissues (Figure 5). Our findings demonstrate that some elusive genes harbor low-level and spatially-restricted expression profiles, which are rarely observed in the non-elusive genes.

**Figure 5.**
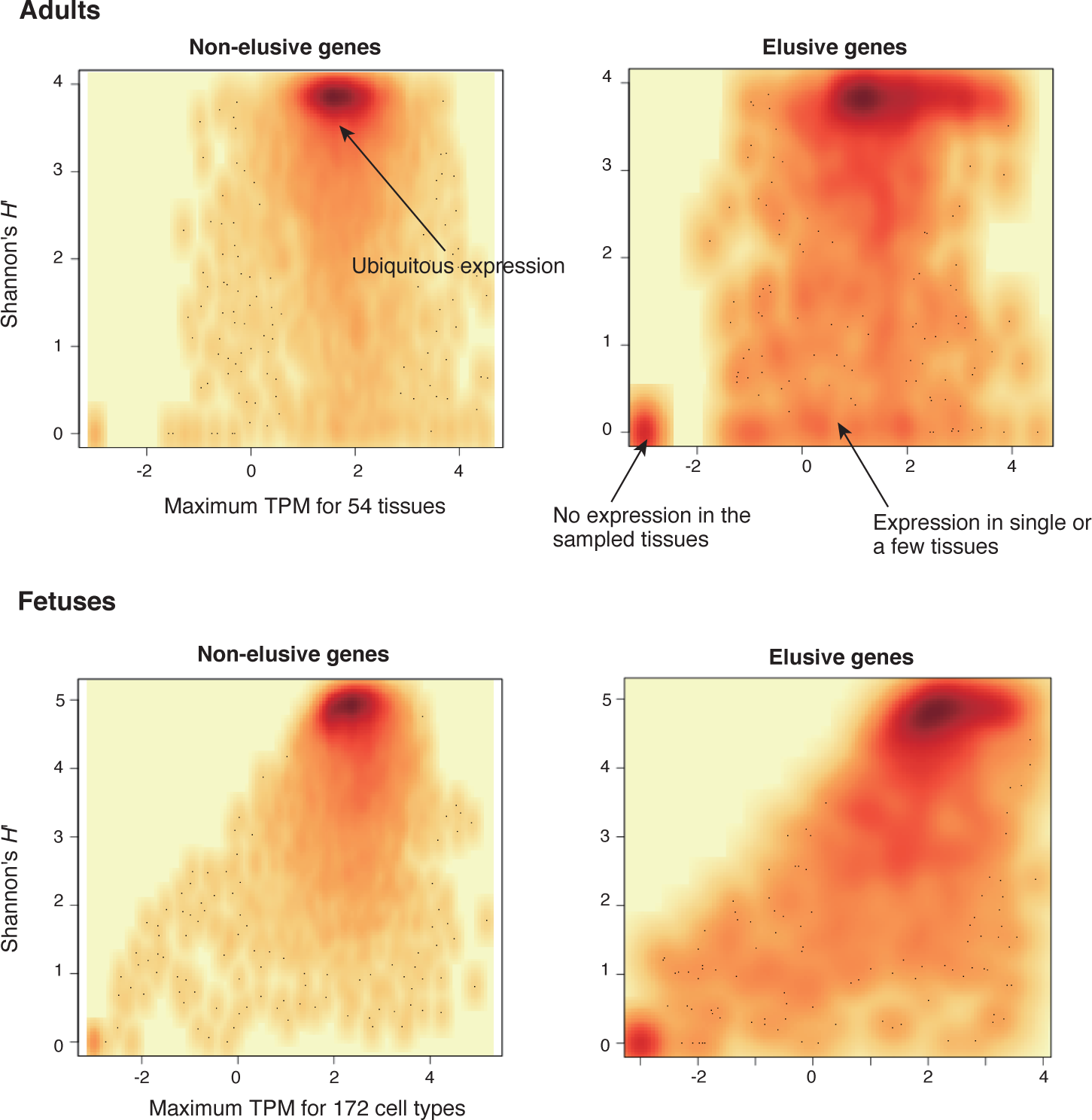
Expression profiles of elusive and non-elusive genes The figure shows density scatter plots of the expression quantity and divergence of elusive and non-elusive genes. The median TPM values of the individual adult tissues across populations were retrieved from the GTEx database (GTEx Consortium, 2020), and normalized TPM values of the fetal cell types were retrieved from the Descartes database (Cao et al., 2020). For the individual genes, maximum TPM and Shannon’s *H*′ values were computed using these processed TPM values.

### Epigenetic nature of elusive genes

Our finding of the low-level and spatially-restricted expression patterns of elusive genes prompted us to explore epigenetic properties involved in this transcriptional regulation. Therefore, we retrieved epigenetic data on a variety of human cell lines from a few regulatory genome databases including ENCODE, a repository that stores the comprehensive annotations of functional elements in the human genome (ENCODE Project Consortium, 2012). Using this information, we characterized the epigenetic features of the genomic regions containing elusive genes (Figure 6).

**Figure 6.**
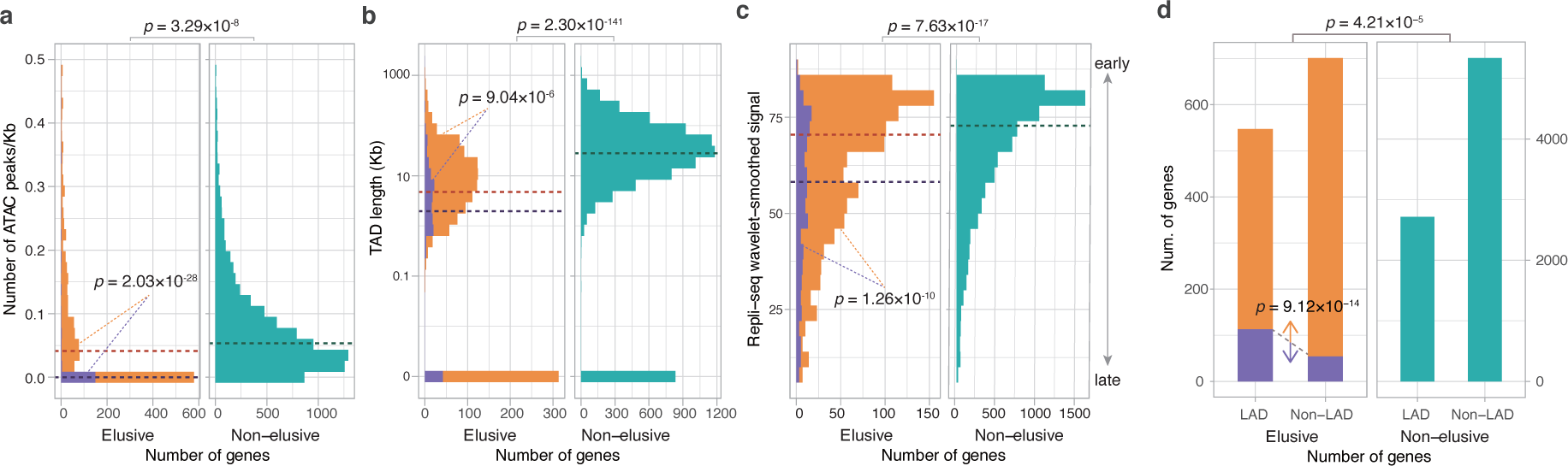
Epigenetic features of the elusive genes Comparison of the distribution of ATAC-seq peak density. (**a**), length of the Topologically Associating Domains (TADs) including the elusive or non-elusive genes (**b**), the replication timing indicator based on Repli-seq (**c**), and overlap with the Lamina-Associated Domains (LADs) computed from Lamin B1 ChIP-seq data. (**d**). ATAC-seq and Hi-C were performed with A549 cells, Repli-seq was performed with HepG2 cells, and Lamin B1 ChIP-seq was performed with HAP-1 cells. In the elusive gene panels, purple and orange bars indicate the elusive genes with restricted expressions (*H*′ < 1; Figure 5) and those with more ubiquitous expressions (*H*′ ≤ 1), respectively. The results for other cells are shown in Supplementary Figures S4–S7. For each epigenetic characteristics, correction for multiple testing was performed for comparison in each cell cultures.

We compared peak densities based on the Assay for Transposase-Accessible Chromatin using sequencing (ATAC-seq), an indicator of accessible chromatin regions in the genome, in the gene bodies and flanking regions between the elusive and non- elusive genes. In seven cell lines out of eight examined (nine experiments of ten), the results showed fewer ATAC-seq peaks in the genomic regions including the elusive genes than in those including non-elusive genes, indicating that the elusive genes are likely to reside in inaccessible genomic regions (Figure 6a; Figure 6–figure supplement 1). We also searched for Topologically Associating Domains (TADs), genomic elements with frequent physical self-interaction potentially acting as promoter-enhancer contacts (Rao et al., 2014) that included either the elusive or non-elusive genes. The result showed that a higher fraction of the elusive genes resided outside of the TADs than the non-elusive genes for all the eleven cell lines investigated (Figure 6b; Figure 6– figure supplement 2). Furthermore, the elusive genes were located in shorter TADs.

These observations suggest that the elusive genes are unlikely to be regulated by distant regulatory elements (Figure 6b).

Our investigations extended to the association of the elusive genes with further global regulation of genomic structures. We compared the percentage normalized signal of Repli-seq (Hansen et al., 2010), a high throughput sequencing for quantifying DNA replication time as a function of genomic position, between the elusive and non-elusive genes. The results showed that elusive genes were prone to late replication in all of the 15 cell lines examined (Figure 6c; Figure 6–figure supplement 1). Late-replicating regions are frequently located at the nuclear periphery and often interact with the nuclear lamina. Therefore, we examined the nuclear position of the genomic regions including the elusive genes by referring to the Lamina Associating Domains (LADs) that were identified by the ChIP-seq reads for Lamin B1 (van Schaik et al., 2020; Zheng et al., 2018). Compared with the non-elusive genes, the elusive genes were found to be enriched in LADs for all of the four cell lines examined (Figure 6d; Figure 6–figure supplement 3), consistent with their late replication timings (van Steensel and Belmont, 2017).

We further investigated the association of the restricted expressions of the elusive genes with epigenetic features. From 988 elusive genes whose expressions were quantified in the GTEx database, we classified the elusive genes into two groups based on the expression diversities: that is, 173 elusive genes with Shannon’s diversity index *H*′ > 1 were ubiquitously expressed, and 815 of those with *H*′ ≤ 1 were expressed in only a few or none of the tissues examined (Figure 5). Importantly, all of the four epigenetic features of the elusive genes with *H*′ ≤ 1 were more pronounced than those with *H*′ > 1: sparse ATAC-seq peaks, short TADs, late replication timings, and significant overlaps with LADs (Figure 6; Figure 6–figure supplement 4). This observation suggests that low-level and spatially-restricted expressions of the elusive genes are associated with intrinsic epigenetic features of these genomic regions.

High GC contents in genomic regions potentially hinder identifying an epigenetic feature by short read sequencing because of underrepresentation of sequence reads by amplification-based sequencing libraries. This bias might lead to sparse distributions of the ATAC-seq peaks and Hi-C contacts in the genomic regions that contain the elusive genes. However, only 8.09% and 10.8% of the elusive genes with *H*′ ≤ 1 and *H*′ > 1 were located in extremely high GC-content regions (>60%), respectively, with no significant difference (*p* = 0.337). Thus, the depleted epigenomic features in the genomic regions containing elusive genes are unlikely to be false discoveries caused by a technical issue, namely, underrepresentation of the sequencing reads.

## Discussion

Here we identified elusive genes that were lost in multiple lineages during mammalian evolution, using a comprehensive scan of gene phylogenies. To identify gene loss events, absence of evidence (i.e., missing genes caused by incomplete genome assemblies and gene annotations), should be reviewed meticulously (Deutekom et al., 2019). Additionally, gene loss might be detected erroneously because of failure in similarity searches for orthologs of rapidly evolving genes (Moyers and Zhang, 2015). In this study, we aimed to reduce these false discoveries through our multifaceted approaches (Figure 1). We selected those species with highly complete gene annotations through integration of multiple gene annotations. Using these improved gene annotations, we created orthologous groups by employing a highly sensitive homology search with MMSeq2 (Steinegger and Söding, 2017) and merged them into those identified in the Ensembl database. Furthermore, we restricted the loss events that were observed as gene absence in all species examined within all hierarchical levels of the selected taxonomic groups (Figure 1b). This absence is likely to have occurred as a gene loss in the common ancestor of the particular taxon rather than as a false discovery of gene loss in the individual species independently. Genuine continuous (e.g., telomere-to-telomere) genome assemblies are now available using modern sequencing technologies (Nurk et al., 2022). These genomic assemblies may help relieve the labor of examining for information losses, thereby facilitating the identification of genuine gene loss in any given species.

In the human genome, the elusive genes and their flanking regions harbor particular characteristics, including high GC-content and high gene density, that may have originated long before the emergence of mammals (Figure 3). Frequent synonymous variations across modern humans in the elusive genes, consistent with higher synonymous substitution rates between the vertebrate orthologs, suggest that the genomic regions including elusive genes have been subject to rapid evolution for 500 million years (Figures 2 and 4). Our findings indicate that heterogeneous genomic characteristics potentially affect the fate of genes at the latest period of vertebrate evolution. Analyses with large numbers of germline mutations in the human genome have illustrated the heterogeneity of mutation rates (Campbell and Eichler, 2013; Seplyarskiy and Sunyaev, 2021; Terekhanova et al., 2017). High GC-content in the elusive genes may have facilitated an elevation of the mutation rate, as observed in the enrichment of rare variants in high-GC regions in the human genome (Schaibley et al., 2013). In addition, some of the elusive genes appear to have retained particular epigenetic marks including sparse ATAC-seq peaks, late replication timings, and location within LADs (Figure 6; Supplementary Figures S4–S7); these epigenetic marks are relevant to an increase in the mutation rate. Genomic regions with late replication timing exhibit increased mutation rates because of their unstable structure during the S-phase of the cell cycle (Koren et al., 2012; Stamatoyannopoulos et al., 2009). LADs retain more G-to-A mutations because of their susceptibility to oxidative damage in the nuclear periphery resulting in high levels of 8-Oxoguanine (Yoshihara et al., 2014).

The epigenetic marks of elusive genes are relevant to the suppression of gene expression (van Steensel and Belmont, 2017), and indeed, these genes harbor weakened and spatially restricted expression profiles (Figures 5–6 and S4–S7). However, the genomic features associated with these epigenetic marks usually exhibit lower GC-contents and reduced gene density (Gilbert et al., 2004; Rao et al., 2014; van Steensel and Belmont, 2017). This discrepancy may be caused in part by a gain of local heterochromatin accompanied with suppression of the expression of transposable elements, as observed in various eukaryotic genomes (Choi and Lee, 2020; Fiston-Lavier et al., 2007; Grewal and Jia, 2007; Rangasamy, 2013; Slotkin and Martienssen, 2007; Underwood et al., 2017). Previous analyses showed frequent heterochromatinization of the human genomic regions where KRAB zinc finger genes colocalize with L1 retrotransposons (Imbeault et al., 2017; O’Geen et al., 2007; Vogel et al., 2006). One of the genomic regions found in human chromosome region 19p12 also contains many elusive genes (Vogel et al., 2006) (Fig. 7). Closer attention to the local gene and repeat contents including repetitive elements and tandem gene clusters might facilitate our understanding of heterochromatinization in restricted genomic regions, although we excluded such gene clusters in our search for elusive genes (Figure 1).

**Figure 7.**
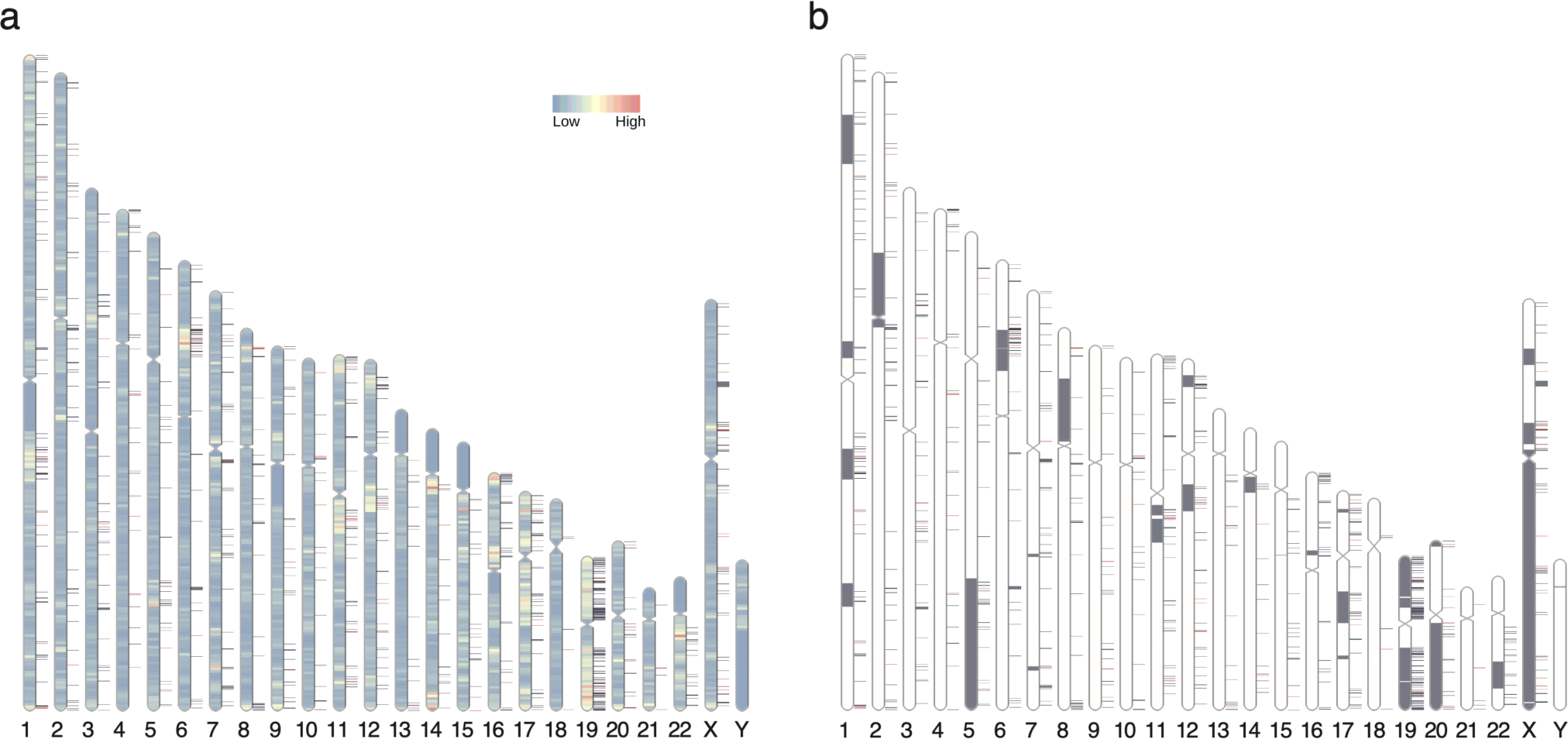
Chromosomal distribution of human elusive genes Red and dark blue horizontal bars on the side of the chromosome diagram represent the location of elusive genes with restricted expression (Shannon’s *H*′ ≤ σ 1) and more ubiquitous expression (*H*′ > 1), respectively. (**a**) The chromosome diagrams are colored according to gene density (number of genes/Mb). (**b**) Gray regions in the diagram indicate orthologous regions of microchromosomes in the ancestors of gnathostomes (Nakatani et al., 2021). The chromosome diagrams were drawn using RIdeogram (Hao et al., 2020).

The heterogeneous locations of elusive genes can also be interpreted from a chromosome-scale viewpoint (Figure 7a; Figure 7–figure supplement 1). Elusive genes were found in particular genomic regions including nearly all of human chromosome 19, and these regions clearly overlapped with regions of high gene density. This is consistent with our observation that the elusive genes were located in the genomic regions with higher gene density than those with the non-elusive genes. Importantly, some of these genomic regions were traced back to the microchromosomes of the ancestral gnathostomes and/or amniotes by karyotyping the ancestral genomes (Figure 7b; Figure 7–figure supplement 1). Recent studies have indicated that these microchromosomes were generated from duplicated chromosomes via allotetraploidization in early vertebrate evolution followed by rapid deletion of large parts of the chromosomal regions (Nakatani et al., 2021; Simakov et al., 2020). In addition, vertebrate microchromosomes harbor particular genomic features including high GC-content, high gene density, and high recombination rate, some of which are concordant with genomic regions containing elusive genes (Groenen et al., 2009; International Chicken Genome Sequencing Consortium, 2004; Schield et al., 2019). This inference of ancestral karyotypes augments our observations that some elusive natures have been retained for hundreds of millions of years, and further suggests that the disparity of genomic regions has been retained for an equivalent timescale.

Finally, we note the potential evolutionary courses that facilitate the transition of gene fate from retention to loss. One possible course is a decrease in essential functions because of rapid sequence evolution in local genomic regions. The elusive genes located in those genomic regions with rapidly-evolving characteristics are likely to accumulate neutral or even moderately harmful mutations in coding regions frequently, resulting in impaired essential functions. Another factor is the spatiotemporal suppression of gene expression via epigenetic constraints. Elusive genes with restricted expressions may have limited opportunities to function, potentially leading to loss of their important roles. The extent of these evolutionary forces may have varied with time and lineages, resulting in patchy loss of elusive genes phylogenetically. Interestingly, a recent large-scale scan of *de novo* mutations in *Arabidopsis* indicates the association of mutation rates with epigenetic features and functional essentiality of genes (Monroe et al., 2022). Further investigation of the association of genes with the surrounding genomic regions in various taxa may provide a common understanding of genomic and epigenomic features that potentially alter the fate of genes. Although epigenetic features are plastic, our findings indicate that the disparities of genomic regions are reflected in the heterogeneity of evolutionary forces and have been retained for hundreds of millions of years. This idea prompts us to explore evolutionary constraints on more global genomic regions that are potentially associated with structural characteristics including chromosomal composition and locations within the nucleus.

## Materials and Methods

### Sequence retrieval

We retrieved genome assemblies and gene annotations of 114 mammals and 132 non-mammal vertebrates from RefSeq (accessed on April 9, 2018), Ensembl release 92, and the repositories of the individual genome projects (Supplementary Table S1). Gene annotations for a single species from multiple repositories were integrated into one as follows. When gene annotations of multiple repositories were generated referring to the same version of the genome assembly, the annotation GTF files were merged with the ‘cuffcompare’ tool (Trapnell et al., 2012). Otherwise, translated amino acid sequences were clustered by CD-HIT v. 4.6.4 (Fu et al., 2012) with 100% sequence similarity, and the representative sequence for each cluster was retrieved by assuming that each cluster represented a single locus. Subsequently, we selected the canonical amino acid sequence for each locus: canonical peptides of the Ensembl genes were retrieved from the Ensembl database; for other resources, the longest amino acid sequence from the isoforms of a locus was chosen. The completeness of the gene annotations was performed on the gVolante web server with assessments by BUSCO v. 2 (Simão et al., 2015) by referring to the CVG (Hara et al., 2015) and BUSCO vertebrate ortholog sets. The gene annotations of mammals, birds, and ray-finned fishes that had fewer than 1% missing genes, as well as those of the other vertebrates with fewer than 3% missing genes, were selected. Exceptionally, the gene annotations of *Gavialis gangeticus* (Reptilia; CVG missing ratio 3.86%), *Paroedura picta* (Reptilia; BUSCO vertebrate ortholog missing rate 3.25%), and *Scyliorhinus torazame* (Chondrichthyes; BUSCO vertebrate ortholog missing rate 4.45%) were added. Finally, the amino acid sequence set of 90 mammals and 101 non-mammalian vertebrates was subjected to t ortholog clustering. We also retrieved coding nucleotide sequences of the canonical amino acid sequences.

### Ortholog clustering and tree inference

We retrieved gene trees of human protein-coding genes and their homologs from Ensembl Gene Tree release 92. From these gene trees, we constructed an amino acid sequence set of the homologs consisting of the species selected in the above section. This sequence set, restricted to Ensembl sequences only, was used as the ‘backbone’ of the ortholog set of all the selected species. In addition, we generated ortholog groups for all the species used by employing OrthoFinder2 v. 2.3.3 (Emms and Kelly, 2019) based on the similarity of amino acid sequences: a sequence similarity search was performed using MMSeqs2 v. 2339462c06eab0bee64e4fc0ebebf7707f6e53fd (Steinegger and Söding, 2017). The Ensembl and OrthoFinder ortholog sets were then merged to create the united set of ortholog groups, yielding 50,768 vertebrate ortholog groups.

The integrated ortholog groups were then subjected to molecular phylogenetic analysis. Amino acid sequences of the individual groups were aligned with MAFFT v. 7.402 (Katoh and Standley, 2013), and ambiguous alignment sites were removed with trimAl v1.4 (Capella-Gutiérrez et al., 2009). Phylogenetic trees were inferred with IQ-Tree v. 1.6.6 (Nguyen et al., 2015) by selecting the optimal amino acid substitution model with ModelFinder (Kalyaanamoorthy et al., 2017) implemented in the IQ-Tree tool for each sequence alignment. In the inferred phylogenetic trees, ambiguously bifurcated nodes—those with branch lengths less than 0.0025—were collapsed into a multifurcational node by the ‘di2multi’ function implemented in ape v. 5.5 (Paradis and Schliep, 2019). The trees were then rooted with the automatic rooting function ‘get_age_balanced_outgroup’ implemented in ete3 v. 3.1.1 (Huerta-Cepas et al., 2016) to minimize any discrepancy of tree topologies with the taxonomic hierarchy of the species included.

### Identification of elusive genes in the human genome

For the individual trees, orthologs of the human genes were detected by the ‘get_my_evol_events’ function in ete3 (Huerta-Cepas et al., 2007). This function inferred gene duplication nodes in the rooted trees, resulting in separation of the trees into 17,495 subtrees of mammalian ortholog groups containing human genes. The ortholog information was referenced to extract the species with no orthologs to humans . This absence was further assessed by the ortholog annotation of human genes in the Ensembl Gene Tree database.

We selected taxonomic groups for the individual mammalian ortholog groups in which the orthologs were missing in all the species examined (Supplementary Table S1). We restricted our study to gene losses that were likely to have occurred in the common ancestor of particular taxonomic groups, rather than those arising from the incompleteness of gene annotations. When a gene was missing in all the taxonomic groups in the same hierarchy, we considered that the gene was lost in the common ancestor of these groups. Finally, we found 1,233 human genes belonging to the ortholog groups that were absent in two or more taxonomic groups and defined them as elusive genes. We further selected 1,081 elusive genes that harbored three or fewer mammalian paralogs for the following analyses. Similarly, we extracted 8,050 human genes whose orthologs were found in all the mammalian species examined and defined them as non-elusive genes. Because these elusive and non-elusive genes were identified in the GRCh38 human genome, we performed the following analyses referring to that assembly.

### Extraction of genomic and molecular evolutionary characteristics

We calculated the GC content of a gene by using its genomic region including introns and untranslated regions (UTRs). To calculate individual gene densities, we extracted genomic regions containing the genes and their flanking three genes at both ends and divided them by seven. The orthologs of the elusive and non-elusive genes were retrieved from the aforementioned gene trees. Amino acid sequence alignment of the human and the ortholog genes was performed using MAFFT. Nucleotide sequence alignments of the coding regions were generated by ‘back-translation’ of the amino acid sequence alignments by trimAl, simultaneously removing ambiguous alignment sites.

By employing coding nucleotide sequence alignments, numbers of synonymous and non-synonymous substitutions per site were computed using PAML v. 4.9a (Yang, 2007).

### Multiomics analysis

Common and rare SNVs of the human populations were retrieved from dbSNP release 147 (Sherry et al., 2001), and human CNVs were obtained from the Database of Genomic Variants (DGV) release 2016-08-31 (MacDonald et al., 2014). The CNVs were classified into duplication and deletion variants, according to the annotation in DGV. The density of these variants in a gene was computed by dividing the number of variants identified in a gene region by its sequence length. Z-scores, indices of the tolerance against mutations, of synonymous, missense, and loss-of-function mutations of the individual genes were retrieved from gnomAD v. 2.1.1 (Karczewski et al., 2021).

Gene expression quantifications of adult and fetal tissues were retrieved from public databases. Expression profiles of adult tissues were obtained from the GTEx database v. 8 (GTEx Consortium, 2020), computed by averaging TPM values across individuals. Expression profiles of fetal tissues were obtained from the Developmental Single Cell Atlas of gene Regulation and Expression (Descartes) portal (Cao et al., 2020), by calculating averaged TPM values of single cells. The maximum TPM values of the individual genes among the tissues were taken as the representative gene expression levels. As a proxy of the spatial diversity of gene expression, Shannon’s species diversity index (*H*′ values) were computed for the individual genes using the following equation:

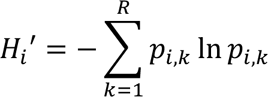

where *Hi*′ represents the Shannon’s index of *i*th gene in the list of the human genes, *pi,k* represents the proportion of the TPM values of the *i*th gene in the *k*th tissues/cell types, and *R* denotes the total number of tissues/cell types examined.

The ATAC-seq peaks and TAD boundaries of the human primary cells and culture strains were retrieved from the ENCODE 3 repository (Accession ID listed in Table S3) (ENCODE Project Consortium, 2012). Wavelet-smoothed signals of the ENCODE Repli-seq data were obtained from the UCSC genome browser (Hansen et al., 2010). The 20 kb bin-associated domains of LAD-seq that employed Lamin B1 antibodies (van Schaik et al., 2020) were retrieved from the 4D Nucleome Data Portal (https://data.4dnucleome.org/publications/f1218a92-1f37-4519-85d6-ccedd5f7ad39).

### Code availability

The scripts for inferring gene presence and absence from gene tree was deposited in GitHub (https://github.com/yuichiroharajpn/ElusiveGenes).

### Statistical tests

Comparisons of the genomic characteristics between the elusive and non-elusive genes were tested statistically with the nonparametric Mann–Whitney *U* test and Fisher’s exact test implemented in R. Correction of multiple testing was performed using the Benjamini–Hochberg false discovery rate (FDR) approach.

## Acknowledgements

We thank Dr. Yoichiro Nakatani for providing the information of orthologous regions of ancestral chromosomes in the human genome. This work was supported by RIKEN to S.K., JSPS KAKENHI Grant Number 20H03269 to S.K. and 21K06132 to Y.H., and Mochida Memorial Foundation for Medical and Pharmaceutical Research to Y.H. Computations were partially performed on the NIG supercomputer at ROIS National Institute of Genetics.

## Supplementary Figures

**Figure 2–figure supplement 1.**
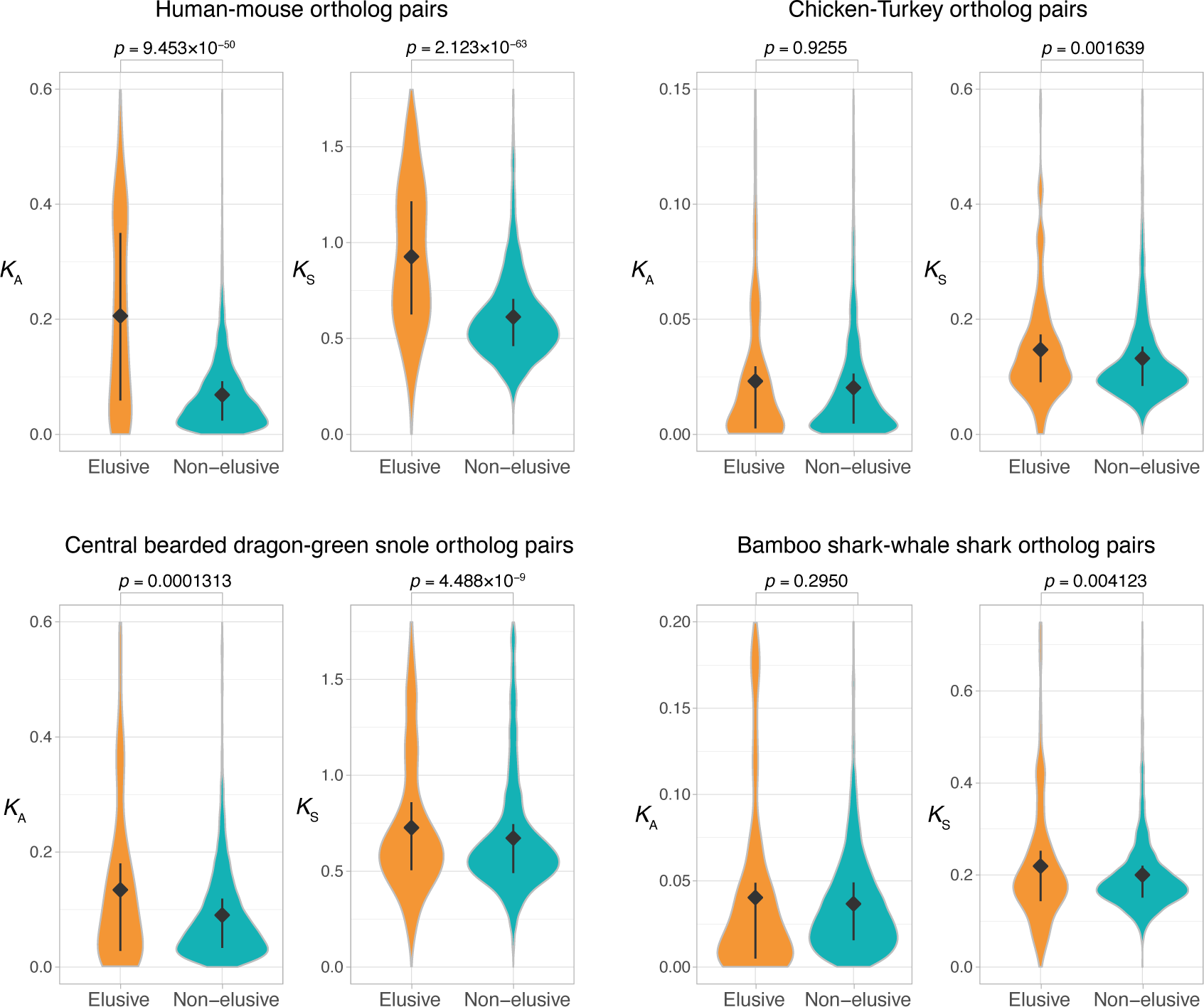
Comparison of *K*_A_ and *K*_S_ values between orthologs of the elusive and non-elusive genes Distributions of *K*_A_ and *K*_S_ values between the orthologs of human elusive and non- elusive genes of closely related vertebrates. Correction for multiple testing was performed for comparison in each species pair.

**Figure 3–figure supplement 1.**
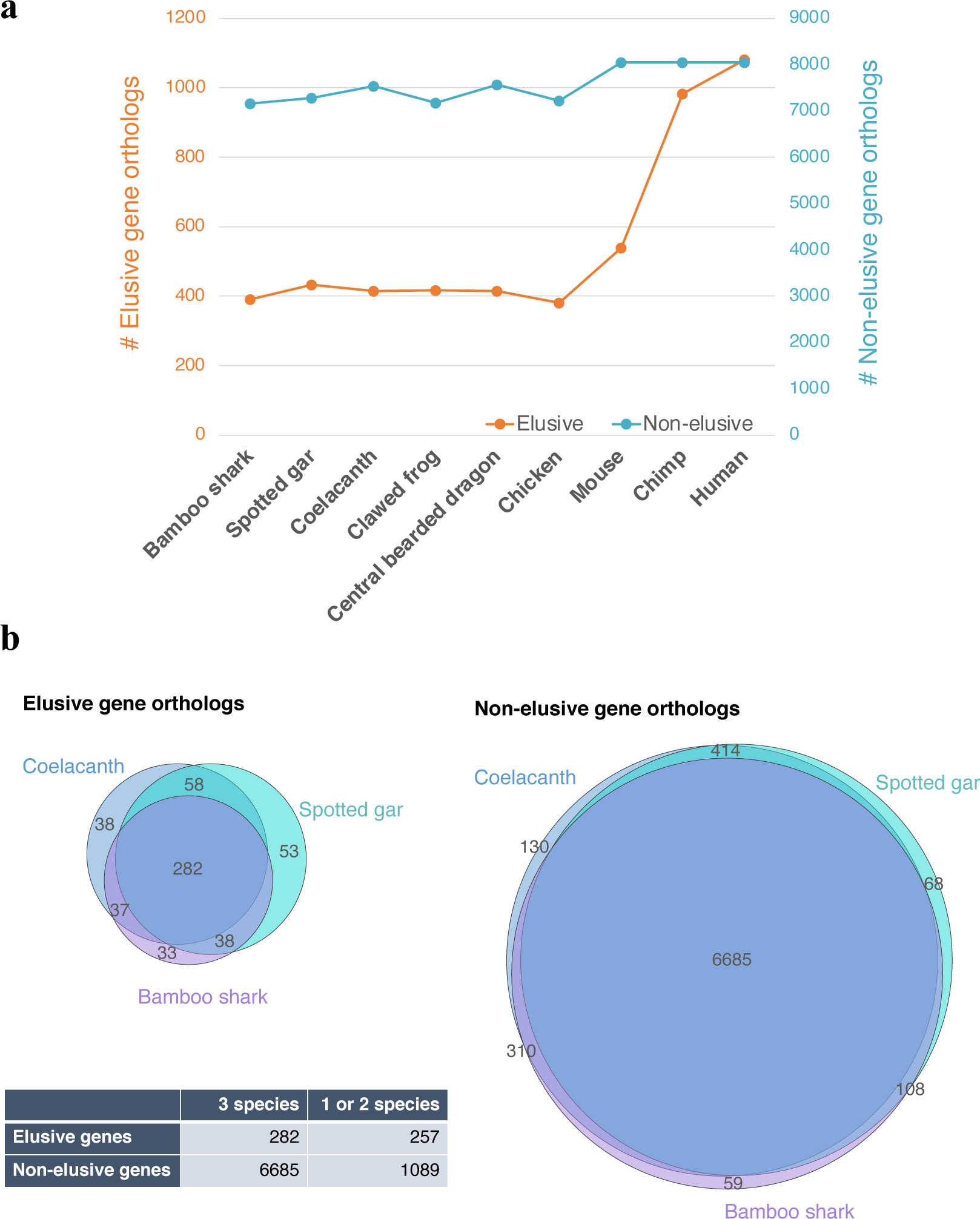
Asymmetric ortholog retention across the vertebrates Number of retained orthologs of the human elusive and non-elusive genes. (**a**) and the overlaps of the retained orthologs across three vertebrates distantly related to modern humans (**b**). The *p*-value of the 2 × 2 contingency table given by Fisher’s exact test is 4.5 × 10^−71^.

**Figure 3–figure supplement 2.**
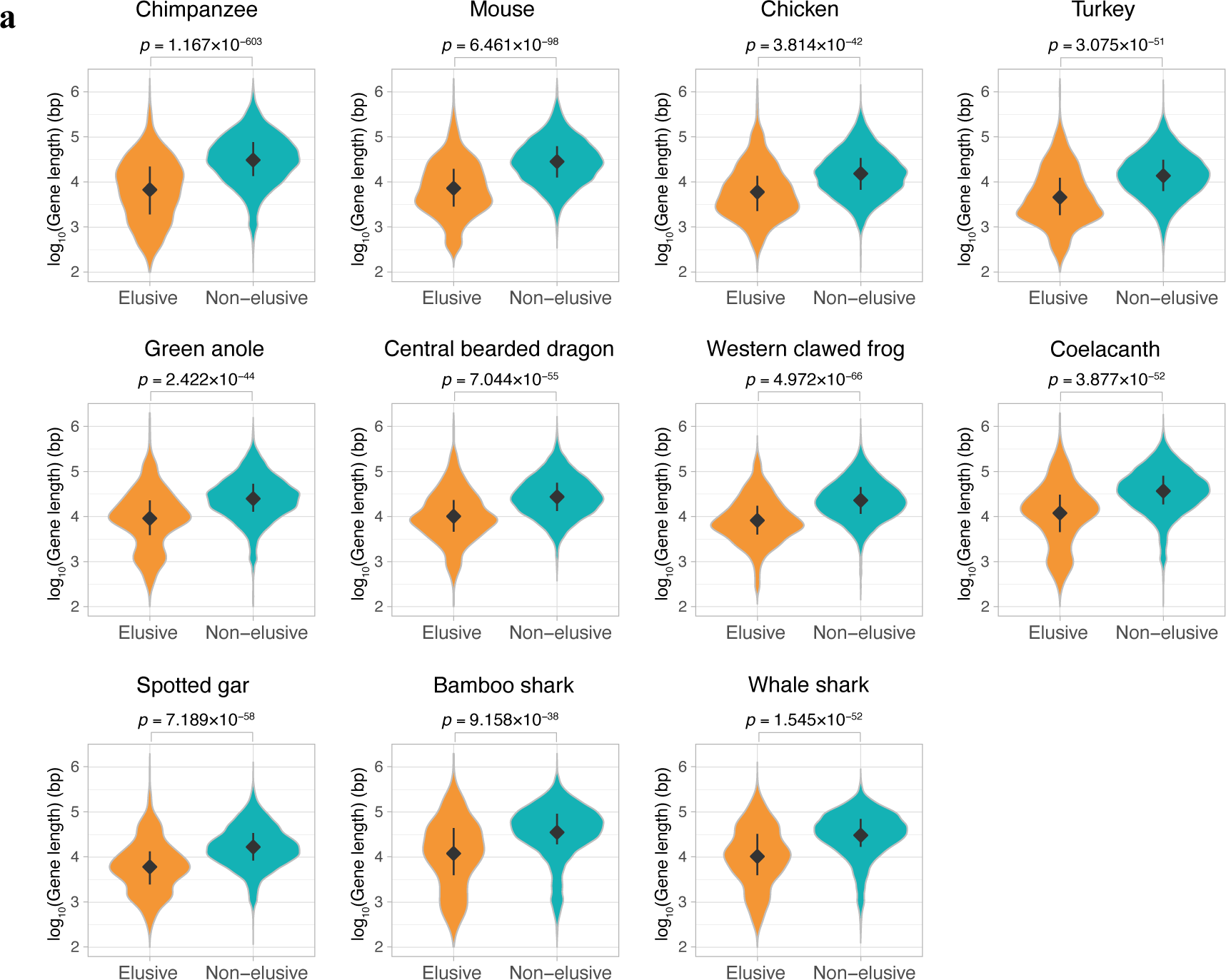
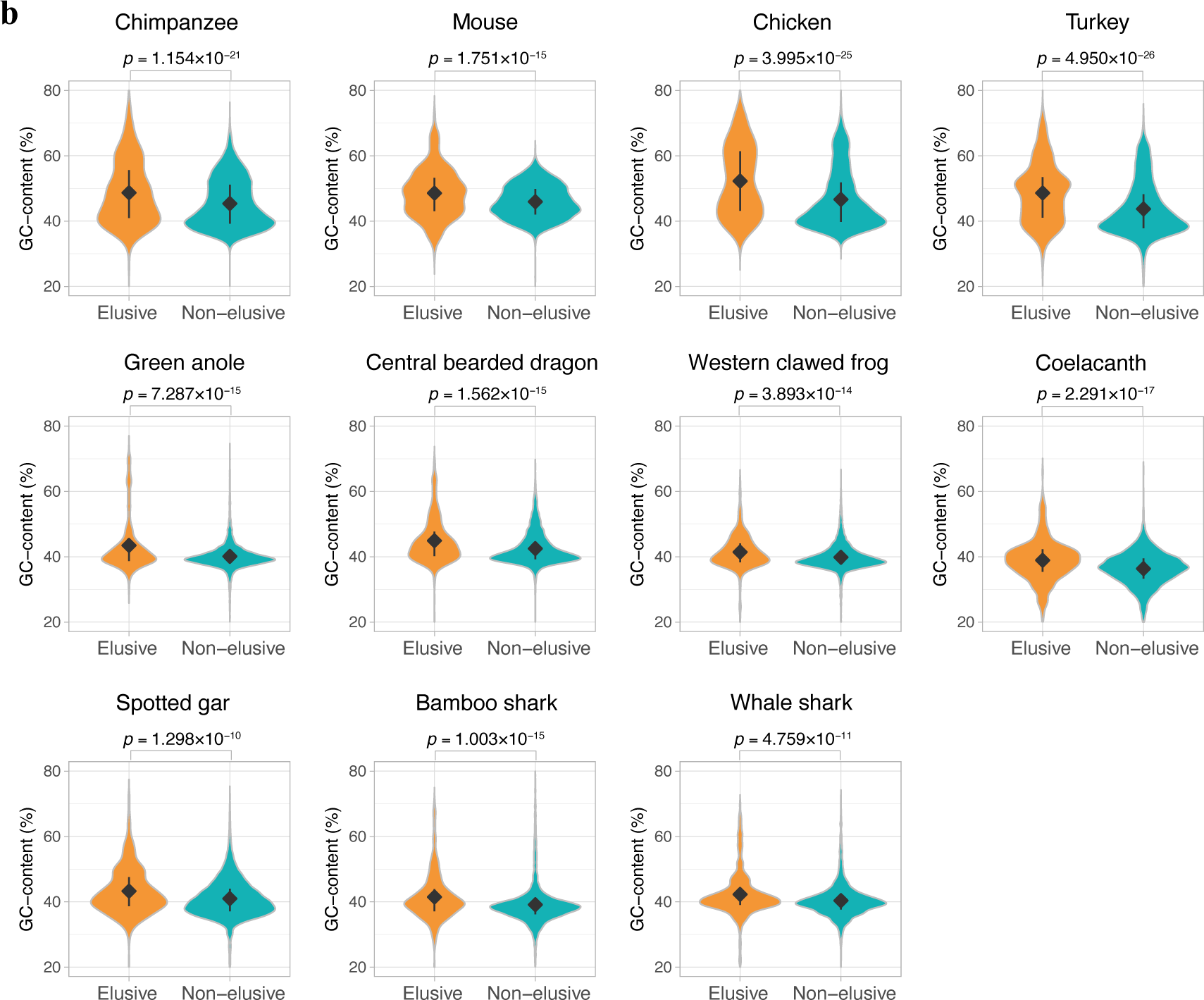
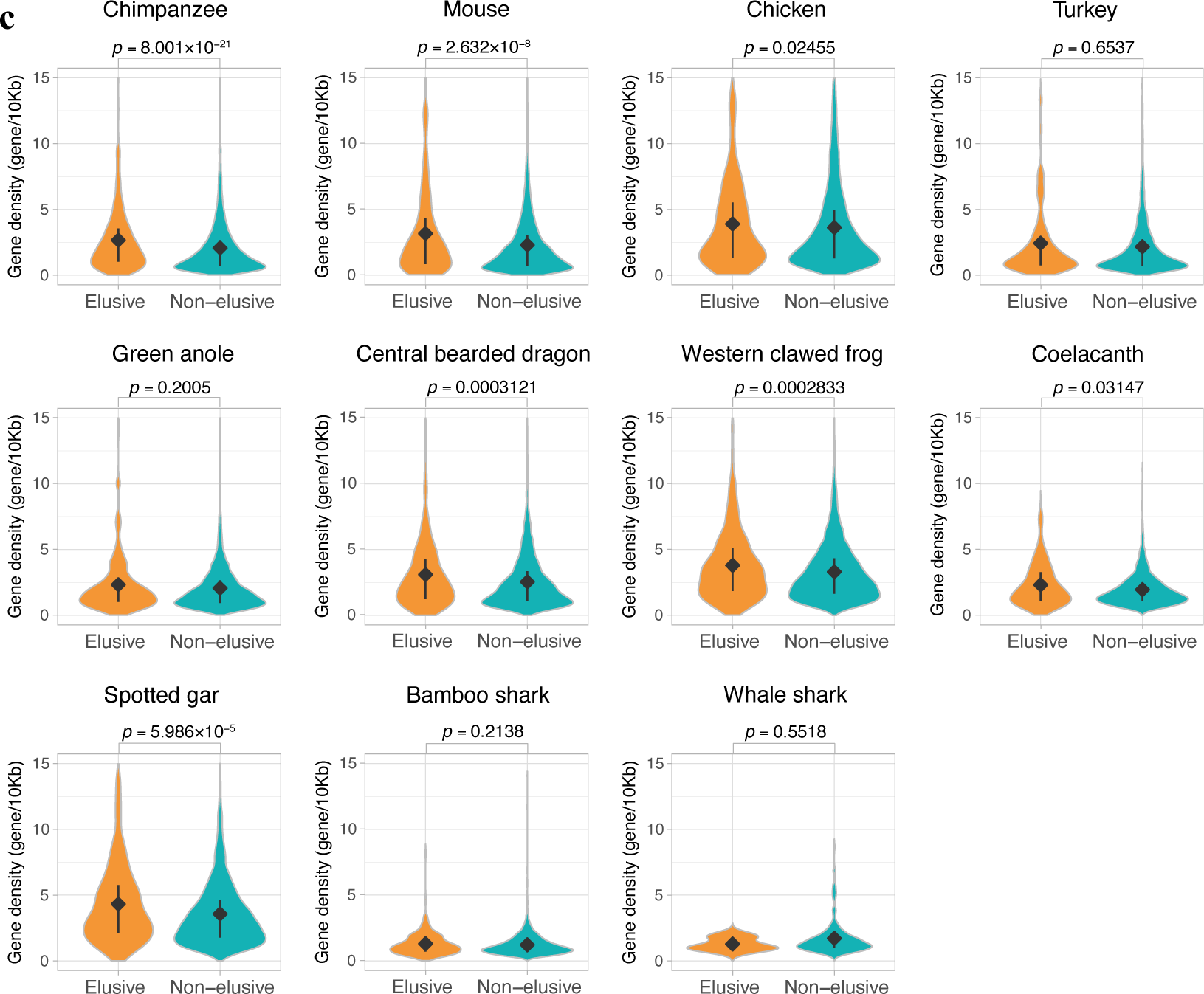
Genomic characteristics of the orthologs of elusive and non-elusive genes Distribution of (**a**) gene length and (**b**) GC-content of the orthologs of the human elusive and non-elusive genes and (**c**) distribution of the gene density of the genomic regions where the orthologs of the human elusive and non-elusive genes are located. For each genomic characteristics, correction for multiple testing was performed for comparison in each species.

**Figure 6–figure supplement 1.**
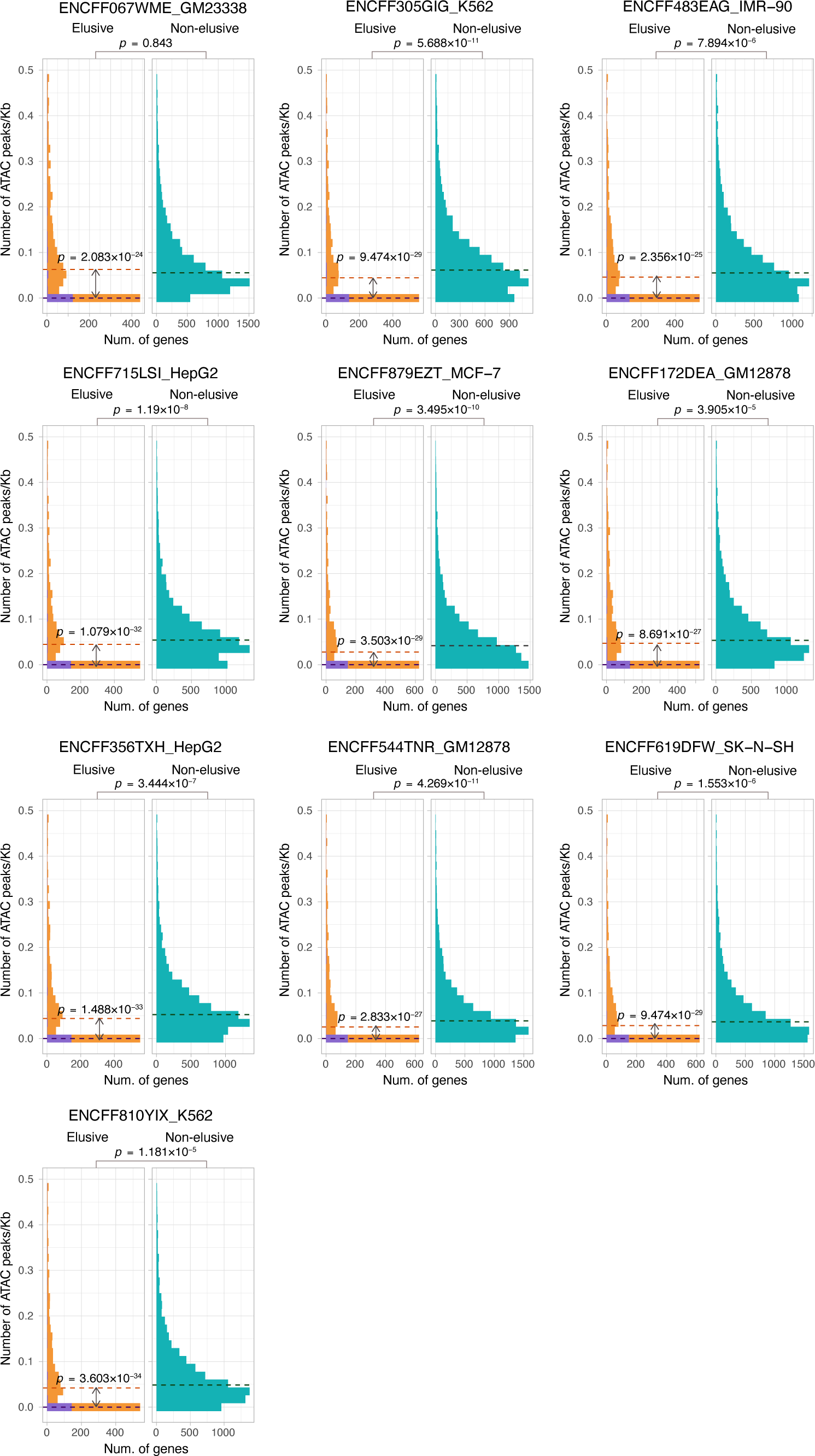
ATAC-seq peak density of the elusive and non- elusive gene regions Comparison of the distribution of ATAC-seq peak density between the elusive and non- elusive genes across multiple cell types. In the elusive gene panels, purple and orange bars indicate the elusive genes with restricted expressions (*H*′ < 1; Figure 5) and those with more ubiquitous expressions (*H*′ ≤ 1), respectively. Correction for multiple testing was performed for comparison in each cell cultures.

**Figure 6–figure supplement 2.**
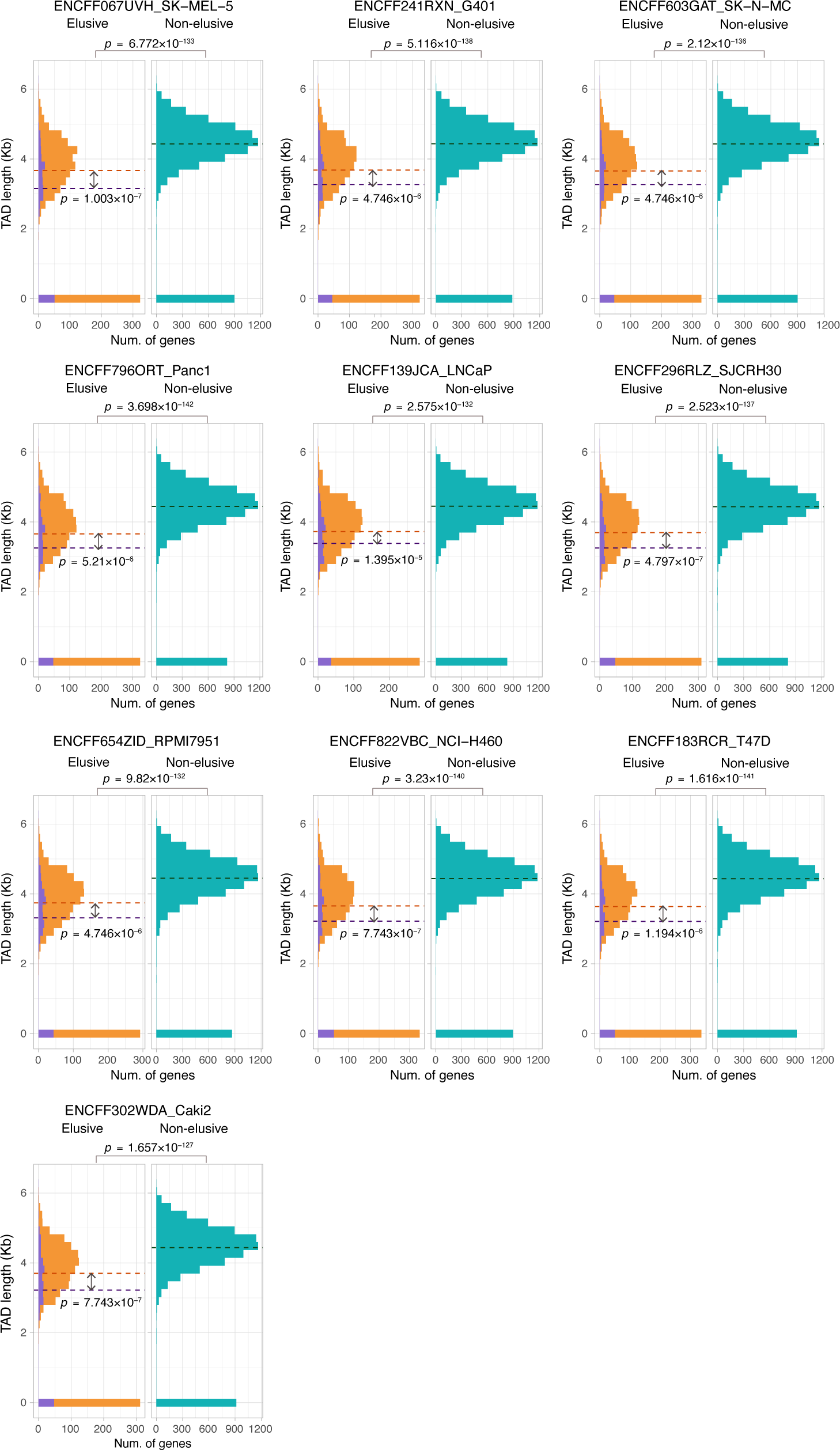
Sequence lengths of the topologically associating domains (TADs) containing elusive or non-elusive genes Comparison of the distribution of length of TADs including the elusive or non-elusive genes across multiple cell types. In the elusive gene panels, purple and orange bars indicate the elusive genes with restricted expressions (*H*′ < 1; Figure 5) and those with more ubiquitous expressions (*H*’ ≤ 1), respectively. Correction for multiple testing was performed for comparison in each cell cultures.

**Figure 6–figure supplement 3.**
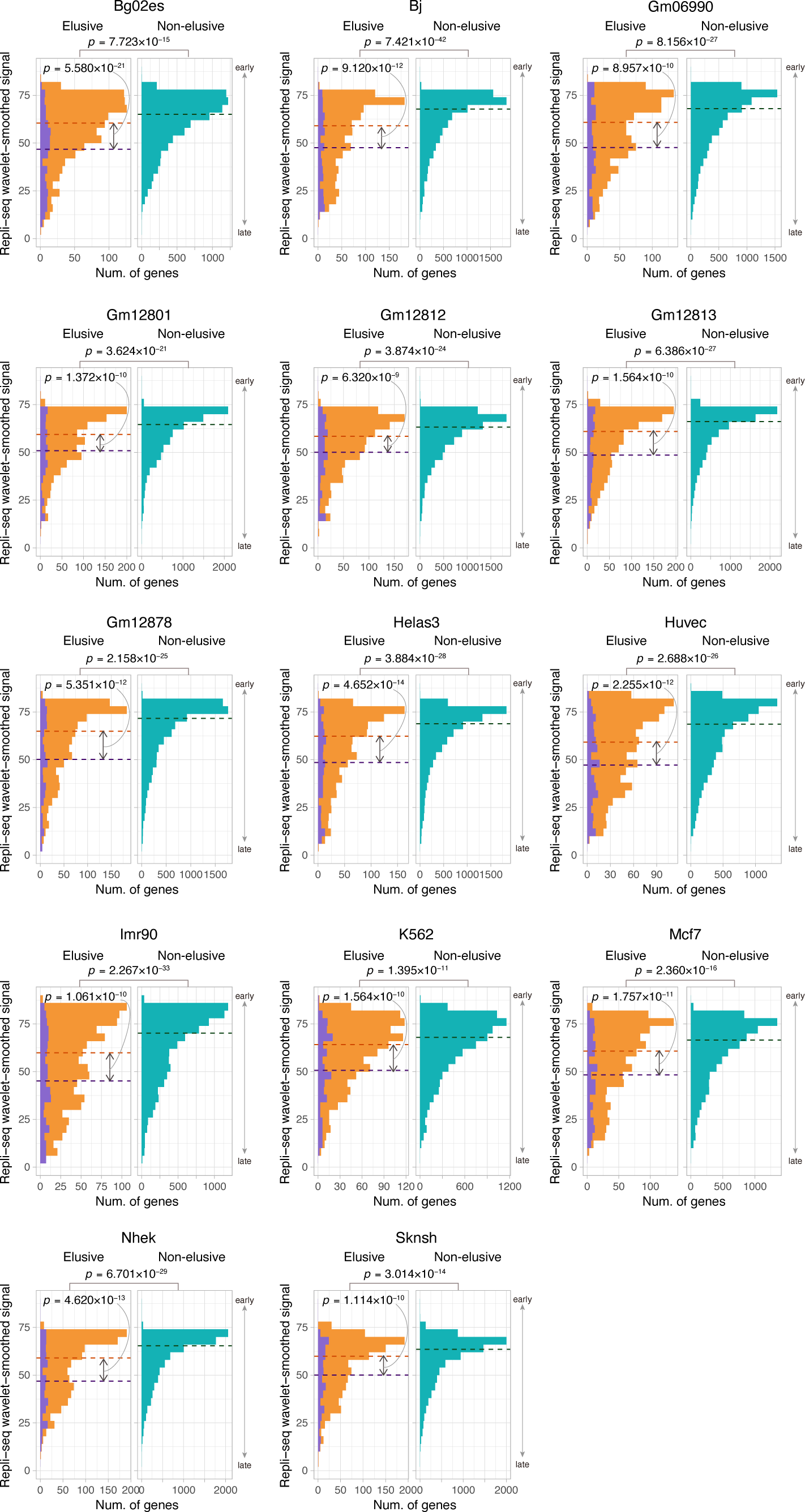
Comparison of the replication timing indicator based on Repli-seq between the elusive and non-elusive genes. Comparison of the distribution of replication timing indicator based on Repli-seq between the elusive and non-elusive genes across multiple cell types. In the elusive gene panels, purple and orange bars indicate elusive genes with restricted expressions (*H*′ < 1; Figure 5) and those with more ubiquitous expressions (*H*′ ≤ 1), respectively. Correction for multiple testing was performed for comparison in each cell cultures.

**Figure 6–figure supplement 4.**
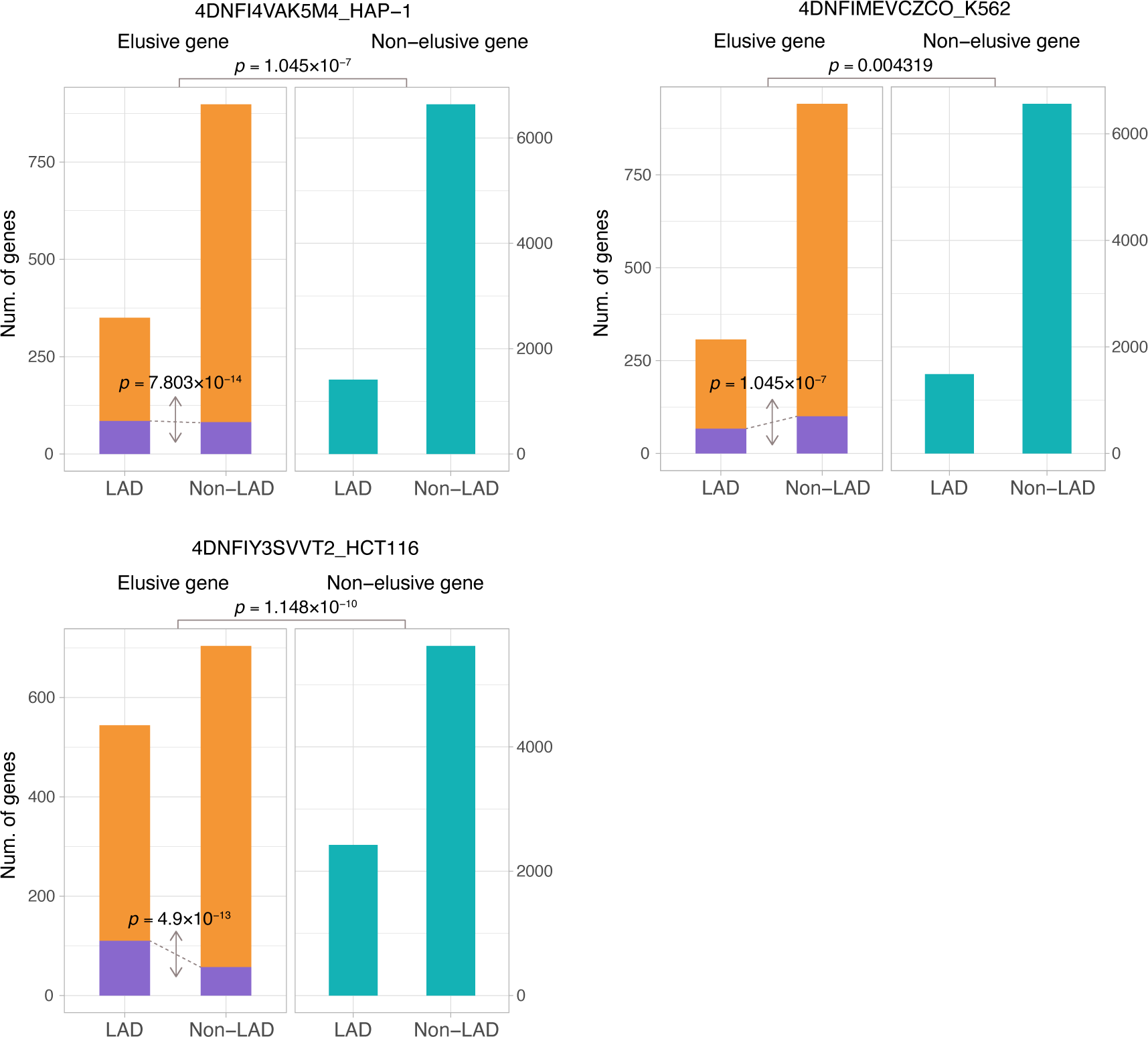
The fraction of elusive and non-elusive genes that overlap with Lamina-Associated Domains (LADs) Comparison of frequency of overlap with LADs computed from Lamin B1 ChIP-seq data between the elusive and non-elusive genes across multiple data. In the elusive gene panels, purple and orange bars indicate elusive genes with restricted expressions (*H*′ < 1; Figure 5) and those with more ubiquitous expressions (*H*′ ≤ 1), respectively. The results for other cells are shown in Figures S4–S7.

**Figure 7–figure supplement 1.**
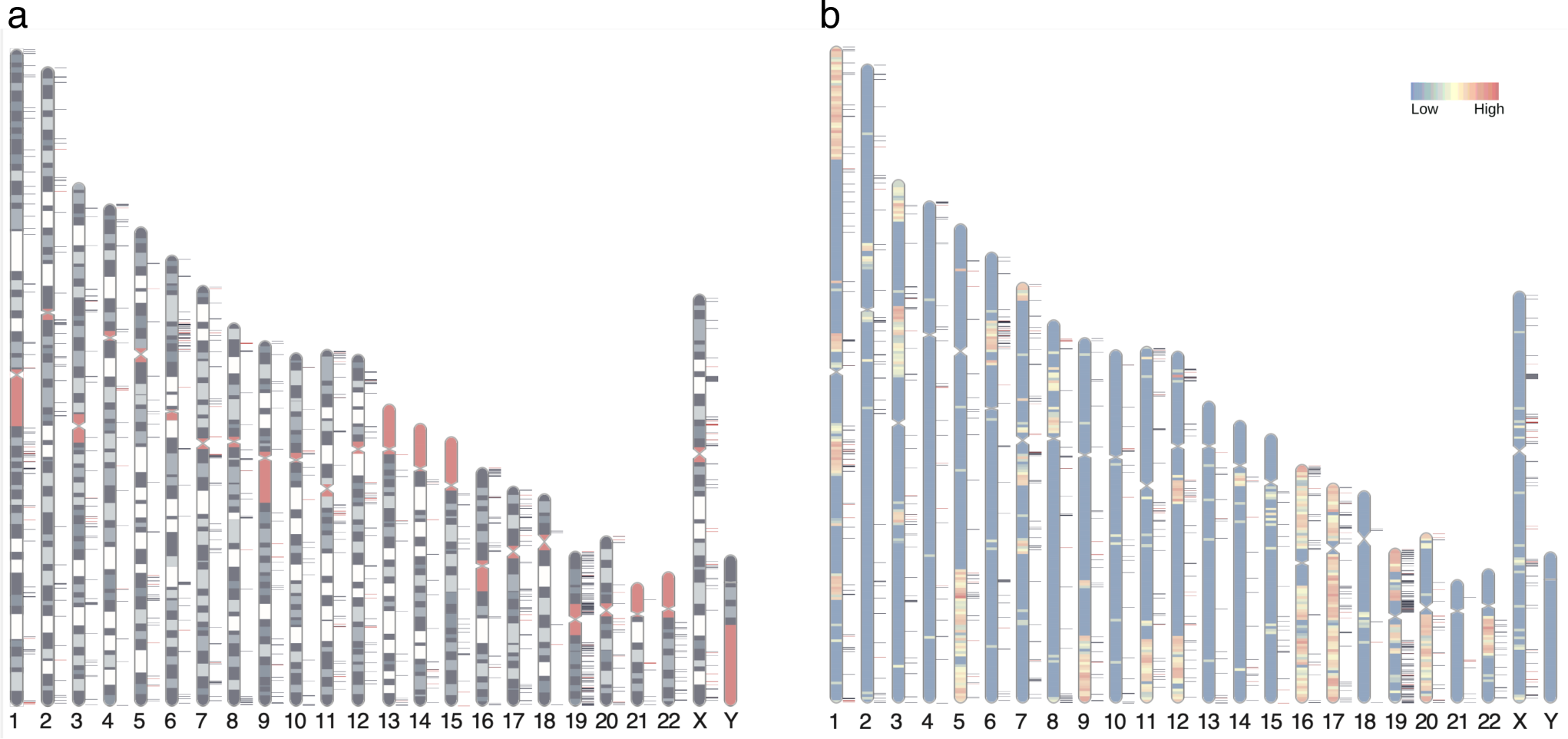
Distribution of elusive genes across human chromosomes Red and dark blue horizontal bars on the side of the chromosome diagram represent the location of elusive genes with restricted expression (Shannon’s *H*′ ≤ 1) and more ubiquitous expression (*H*′ > 1), respectively. (**a**) Karyotypes are shown by G-banding. Red regions indicate centromeres, acrocentric regions, and variable-length regions. (**b**) The chromosome diagrams are colored according to the density of the genes that harbor chicken orthologs in microchromosomes (number of genes/Mb). The chromosome diagrams were drawn using RIdeogram (GTEx Consortium, 2020).

